# Panoramic Insights into the Microevolution and Macroevolution of *Prevotella copri*-containing Lineage in Primate Guts

**DOI:** 10.1101/2020.07.27.224261

**Authors:** Hao Li, Jan P. Meier-Kolthoff, Can-Xin Hu, Zhong-Jie Wang, Jun Zhu, Wei Zheng, Yun Tian, Feng Guo

## Abstract

*Prevotella copri* and related taxa are widely detected in mammalian gut microbiomes and have been linked with one human enterotype. However, their microevolution and macroevolution among hosts are poorly characterized. In this study, extensively collected marker genes and genomes were analyzed to trace their evolutionary history, host specificity, and biogeographic distribution. Investigations based on 16S rRNA gene, *gyrB*, and genomes suggested that a multi-specific *P. copri*-containing lineage (PCL) harbors diverse species in higher primates. Firstly, *P. copri* is the dominant species of PCL in the human gut and consists of multiple groups exhibiting high genomic divergence and conspicuous but non-strict biogeographic pattern. Most African strains with high genomic divergence from other strains were phylogenetically placed near the species root, indicating the co-evolutionary history of *P. copri* and *Homo sapiens*. Secondly, although long-term co-evolution between PCL and higher primates was revealed, sporadic signals of co-speciation and extensive host jumping of PCL members were observed among higher primates. Metagenomic and phylogenetic analyses indicated that *P. copri* and other PCL species found in captive mammals have been recently transmitted from humans. Thirdly, strong evidence was found on the extensively horizontal transfer of genes (e.g., carbohydrate-active enzyme encoding genes) among sympatric *P. copri* groups and PCL species in the same primate host. Our study provides panoramic insights into the complex effects of vertical and horizontal transmission, and potential niche adaption on speciation, host, and biogeographical distribution spanning microevolutionary and macroevolutionary history for a certain gut bacterial lineage.

**Importance:** *Prevotella copri* and its related taxa, which we designated as *Prevotella copri*-containing lineage (PCL) in the present study, are widely detected in guts of human, non-human primates and many captive mammals, showing positive or negative correlation to some human diseases. However, a comprehensive understanding on its microevolutionary (within *P. copri*) and macroevolutionary (among PCL members) history across host species and host biogeography is still lacking. According to our analysis based on 16S rRNA gene, *gyrB* and genomes, we provided the panoramic insights into the putative effects of vertical transfer, horizontal transmission and potential niche selection on host and biogeographical distribution of this gut bacterial lineage and *P. copri*. To our knowledge, it is the first time that a gut bacterial lineage was studied at both micro- and macroevolutionary levels, which can aid our systematic understanding on the host-microbe co-evolutionary interactions.

## Introduction

Animal guts harbor complex microbe assemblies that play key roles in host development, metabolism, and immunity (1–3). Phylosymbiosis between host and gut microbiome has been widely investigated at the community level, and many microbial assemblies show congruence with their host phylogeny (4–6). However, such congruence may not necessarily result from long-term co-evolution between hosts and symbionts (i.e., vertical transfer, which includes transfer along the lineage of multiple host species or among conspecific hosts); other factors, such as diet, host physiology, and immunology, etc. may play uncharacterized roles in shaping gut microbiomes and host phylogeny (7, 8). Alternatively, focusing on certain microbial lineages is a reliable and direct way to trace the history of vertical transfer (9–12). Co-evolution of a bacterial lineage within a set of hosts can be viewed at two levels, namely, macroevolution (interspecies) and microevolution (intraspecies). The former is often discovered among remotely related host species (9, 10, 13, 14), and the latter is observed on hosts belonging to either the same or different host species (12, 15).

In addition to vertical transfer, the biogeographical and host-specific distribution of certain gut bacterial lineages could be largely influenced by the potential horizontal transfer among heterospecific hosts and ecological selection (16–18). Allochthonous taxa may switch to new hosts and initiate new evolutionary branches, potentially causing promiscuous generalists in many host species (19, 20). Novel host-microbe interactions and adaptations can be introduced in such a scenario (21). However, only few gut bacterial lineages have been comprehensively studied at the microevolutionary and macroevolutionary levels to understand the potential effects of vertical and horizontal transfers on their biogeography and host-specificity. This limitation is caused by the lack of comprehensive information for a certain lineage from a wide range of host species and geographic regions and the potential extinction of hosts and microbes. In addition, samples must be collected from wild animals barely affected by humans as a prerequisite to minimize artificial interferences (22, 23). However, data from wild animals are usually less available than those from captive animals.

*Prevotella* is a representative genus of one human enterotype (24, 25). Multiple species in this genus have been detected in human feces. *Prevotella copri* and a few closely related species have the highest frequency and abundance (26) and have been linked with a few human diseases as potential disadvantageous factors (27, 28). On the one hand, two studies based on single-nucleotide polymorphisms have preliminarily reported intraspecies diversification, biogeography, and microevolution (15, 29). A recent work collected over 1,000 *P. copri*-related genome bins using a reference-based metagenomic binning strategy and reported four species-level clades occurring in human guts, thereby validating the species-level diversification in this lineage (30). However, phylogeny conducted solely based on genome bins may lose the overall diversity from minor and rare organisms that could not be genomically retrieved from metagenomes. On the other hand, *P. copri* and related taxa have been frequently discovered in the gut microbiomes of nonhuman primates and other mammals (31–33). Panoramic insights into the macroevolutionary and microevolutionary history of *P. copri* related with its host phylogeny across mammals, primates, and humans remain unclear.

In this study, we reconstructed the robust phylogeny of *P. copri* and its related taxa by comprehensively collecting their phylogenetic markers from multiple hosts and various geographic regions. The rRNA gene, gene encoding DNA gyrase subunit B (*gyrB*) sequences, and genomes were used as a reference to solve interspecies phylogeny, which showed the existence of a multi-specific *P. copri*-containing lineage (PCL). The genomes and a selected marker gene were employed as a basis in investigating the intraspecific phylogeny and microevolution of *P. copri*. With regard to the phylogenies at different levels, vertical and horizontal transmission of *P. copri* and related species can be deduced.

## Results

### Existence of a *P. copri*-containing lineage based on 16S rRNA gene sequence analysis

Figure 1A shows that among the 36 clone sequences, the type strain DSM 18205 and six isolates formed a lineage within *Prevotella* with moderately supportive BS values. All 43 sequences were obtained from the fecal samples of humans, nonhuman primates, bovines, and humanized mice. Several clone sequences from nonhuman primates were located at the root of the clade and exhibited a deeply branching feature.

**Figure 1.**
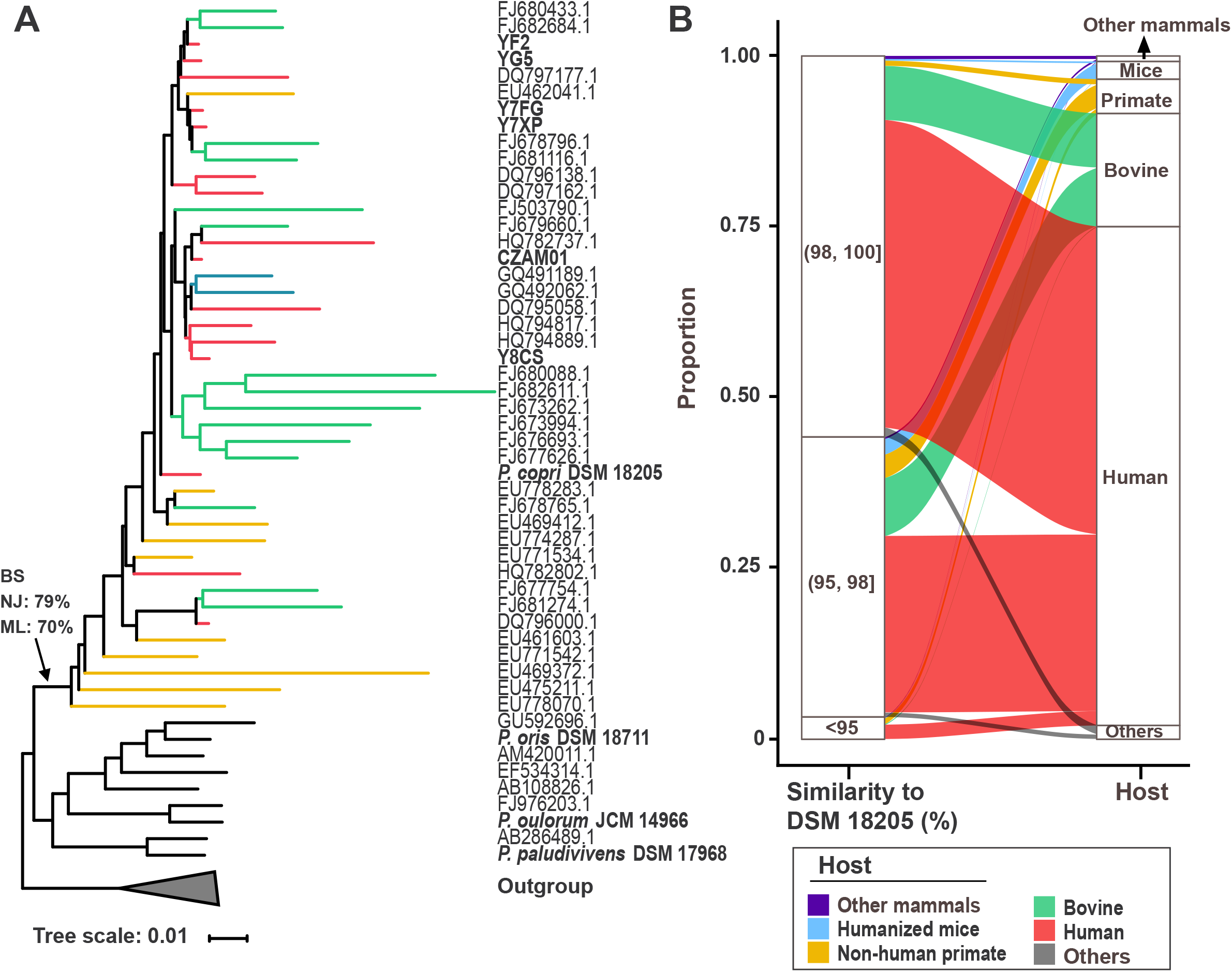
16S rRNA gene-based analysis on the distribution and phylogeny of PCL. A) Neighbor-joining tree based on 534 16S rRNA gene sequences of the genus *Prevotella* with 100 iterations of bootstrapping. Bold labels indicate sequences from isolates. B) Sankey diagram showing the distribution of host origins and similarity fraction (to DSM 18205) for 16S rRNA gene sequences from SILVA database identified as the PCL (*n*=5,316).

A total of 5,316 SILVA 16S rRNA gene sequences putative affiliated with the lineage were retrieved to comprehensively profile the source of this lineage. As shown in Figure 1B, most of the sequences were obtained from the fecal samples of the four hosts, and the rest (<3%) were obtained from other mammals (mostly captive ones such as pig, dog, and mammals from zoos) and human-related environments (e.g., human skin and wastewater). Nonhuman primate-derived sequences contributed over 25% of the remotely related fraction (<95% similarity to the 16S rRNA gene of DSM 18205, Figure 1B) but only accounted for 5% of total sequences. This analysis provides preliminary evidence of a multi-specific PCL, and the results point to its potential macroevolutionary history with primates and the occurrence of PCL members in the guts of captive mammals.

### Phylogeographical pattern of *P. copri* suggests a co-evolutionary history with *Homo sapiens*

A total of 130 *P. copri* genomes (all >70% completeness and <10% contamination, DATA SET S1: Table S1) were used in the phylogenomic reconstruction to investigate the intra-species phylogeny of the most dominant human gut PCL member, *P. copri*. Among which, 116 genomes reached the high-quality criterion (>90% completeness and <5% contamination, according to Bowers et al. (34)). NMPZ01, Y7XP, and Y7FG (belong to the same species) are not *P. copri* according to the average nucleotide identity (ANI) and digital DNA–DNA hybridization (dDDH) values (DATA SET S1: Table S2) and thus were set as the outgroup. The phylogenetic trees reconstructed by concatenating the sequences of 1,095 core single-copy genes and dDDH were highly consistent in terms of topology (Figure 2A). In the core-gene based phylogenetic tree (Figure 2A), the genomes were divided into nine groups (non-monophyletic g0 and monophyletic g1–g8) and three single branches. The six major groups were g0 (*n*=18), g1 (*n*=12), g3 (*n*=9), g4 (*n*=26), g5 (*n*=6), g6 (*n*=43), and g7 (*n*=9) and exhibited well-supported phylogeographical pattern to a great extent. Group g0 was mostly contributed by Africans (only one from USA), g1 and g7 almost exclusively occurred in China (two g1 strains are from Africa), and g5 was only covered by the strains found in Salvador. Groups g4 and g6 consisted of strains from multiple continents, mostly from European countries, USA, and Kazakhstan and a few from China and African countries. Type strain DSM 18205 formed g8 together with one strain from Denmark. Under the dDDH-based species clustering, 130 *P. copri* genomes can be assigned into nine maximally supported, monophyletic species-level clusters or single branches (Figure 2A) (35). Therein, g0 comprises eight clusters, and all the other groups constitute a single cluster. According to both approaches, no close relatives was observed for any of the 130 genome pairs, except for those obtained from the sequential samples of one person (O2.UC17-0 and O2.UC17-1 in g4 and O2.UC38-0 and O2.UC38-2 in g6) and four from a family (M04.1-V3, M04.3-V1, M04.4-V3, and M04.5-V3 in g4).

**Figure 2.**
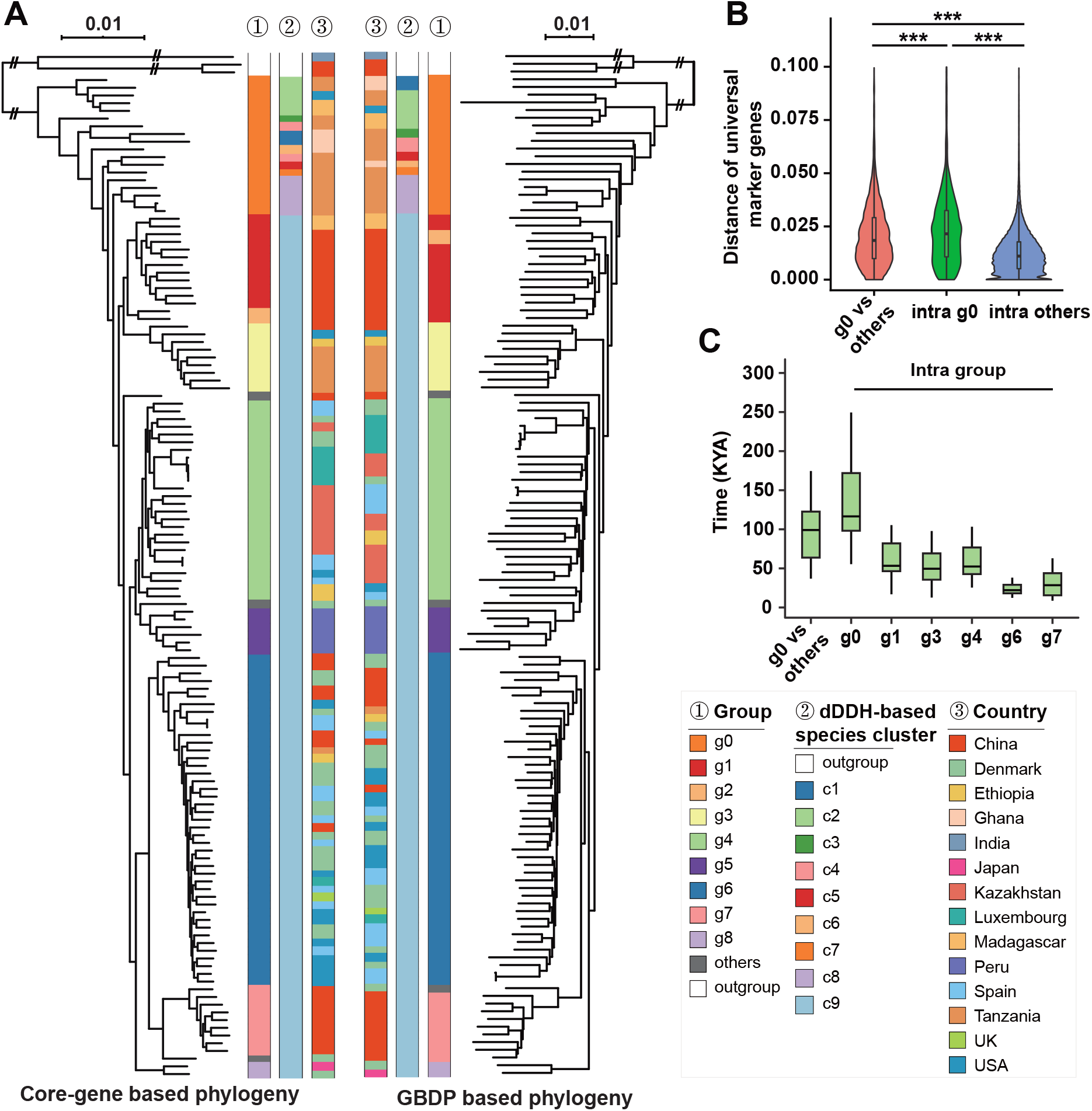
Phylogenomic analyses and molecular dating of 130 globally collected strains primarily belonging to *P. copri*. A) Maximum-likelihood tree of concatenated 1,095 core single-copy orthologous genes (left) and GBDP phylogenomic analysis of the nucleotide sequences restricted to coding regions (right). Values above branches in both trees represent (pseudo) bootstrap support above 60%. Shared annotations include geographical origin ①, clustered groups ②, and dDDH-based species clusters ③. B) Distance of 120 universal marker genes of between group g0 with other strains and within group g0 and other strains. C) Split time of intergroup (g0 to other groups) and intragroup strains.

The co-evolutionary history between the species and *Homo sapiens* was supported by that most strains from Africa were located at the root of the tree and remotely separated from other groups. The large phylogenetic distance between g0 strains and other groups or among g0 strains was further confirmed for single housekeeping genes (Figure 2B). The median synonymous mutation rate of 18 housekeeping genes (without significant intragenic recombination in Pairwise-homoplasy index (PHI) test) from African-derived strains and other strains was 0.052 (DATA SET S1: Table S3). On the basis of the mutation rate of 2.6×10^−7^ per site per year for housekeeping genes of another human symbiont, *Helicobacter pylori* (36), the split of g0 strains and other strains was dated at approximately 99,000 years ago (Figure 2C). This period roughly coincided with the time for modern humans outside Africa [69] and was also supported by *H. pylori* results (37, 38). The median split time for strains for each major group g0, g1, g3, g4, g6, and g7 was 117,000, 53,000, 50,000, 52,000, 22,000 and 28,000 years ago, respectively (Figure 2C).

### Divergence and potential sympatric gene transfer for carbohydrate-active enzymes (CAZys) among *P. copri* groups

*P. copri* was thought to be positively selected by non-westernized diet with high plant-sourced polysaccharides (30, 39, 40). Therefore, the divergence of CAZy modules in the major groups was examined. Pan genomes of the six major groups contained 144 CAZys, nearly half of which were generally distributed in all genomes without significant difference between any two groups (Fisher’s exact test, FDR-corrected *P* >0.05). However, 30 group-specific and 43 sporadic CAZy modules were determined (Figure 3A). High group-specificity was found in several putative alginases (genes containing PL6, PL6_1, and PL17) in g1 and a putative hyaluronidase (the gene containing GH84) in g4 and g6. A gene almost exclusively detected in g1 was identified as a putative alginase by Pfam annotation (PF05426, not annotated against the CAZy database) and functionally verified via the heterologous expression in *Escherichia coli* (cloned from YF2) and biochemical assays (TEXT S1: Figure S1 and SI methods).

**Figure 3.**
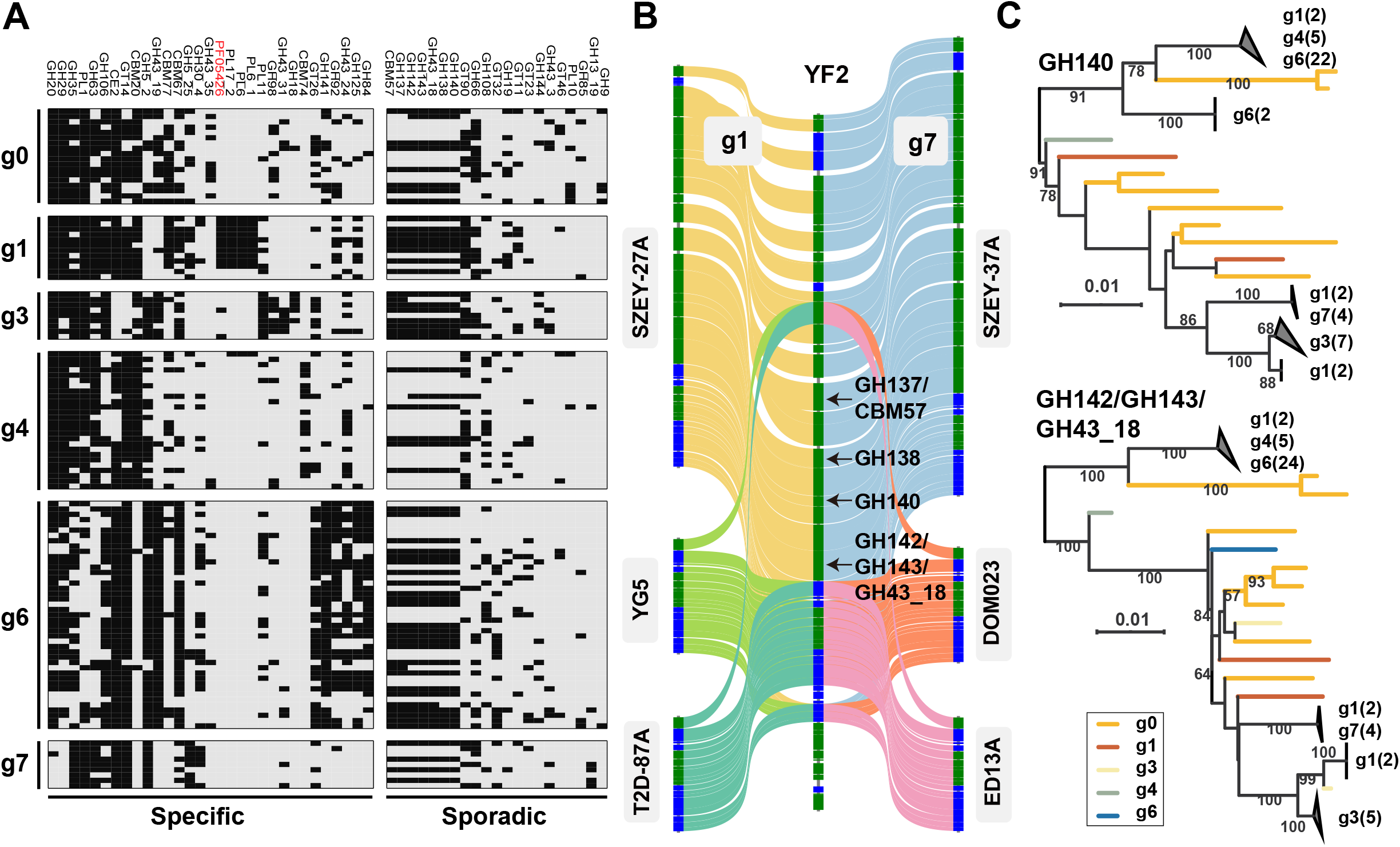
CAZys of the six major *P. copri* groups and their biogeography related microevolution. A) Group-specific (FDR-corrected *P* <0.05 for Fisher’s exact test on the frequency of any groups) or sporadic (detected in less than half of the genomes and no significant difference between any two groups) CAZy families in six major groups are shown in the heatmap (black: present; grey: absent). The gene indicated by the arrow was identified by Pfam annotation using HMMER and functionally verified (TEXT S1: Figure S1). B) Genomic synteny of the fragment containing four sporadic CAZy-encoding genes with flanking genes are displayed for six genomes belonging to g1 (*n*=3) and g7 (*n*=3) using the complete genome of YF2 for mapping. Gene in sense strand (green) or reverse strand (blue) are shown by block colors. C) Maximum-likelihood trees of two sporadic CAZy-encoding genes (in nucleotide) with 100 bootstrap iterations. Sequences exhibiting extremely high similarity (>99%) are collapsed, and their composition is shown.

A sporadic CAZy (containing three modules of GH142, GH143, and GH43_18) was related with a novel depolymerase from *Bacteroides thetaiotaomicron* targeting on complex glycans (>60% amino acid similarity with protein BT1020 (41)). Genomic synteny showed that the absence or presence of the gene-related cluster (containing four CAZy encoding genes) was not due to incorrect assembly or binning (Figure 3B). The gene cluster (approximately 40 kb in length, containing 15 genes in the complete genome of YF2) was colocalized on the same genomic region in all positive strains but was clearly deleted in negative strains. Phylogenies of the two sporadic CAZy-encoding genes located in the cluster revealed inter-group horizontal gene transfer (HGT) for sympatric groups (Figures 3C and S2, the other two CAZy-encoding genes have similar signals but not shown). For example, the gene containing GH140 had three phylogenetic clusters consisting of highly similar sequences (>99.5% amongst all the nucleotide sequences) and strains from geographically co-occurring groups (e.g., g1/g7 in China and g4/g6 mostly in European countries and USA). Investigation on the *P. copri* genomes of g1 (*n*=8), g6 (*n*=1), and g7 (*n*=3) isolates provided by a recently study on the Chinese population to exclude the possibility of genomic contaminations for metagenomic bins (42). The findings further supported that the above phylogenetic crosslinks were not derived from genomic contamination (TEXT S1: Figure S2).

### Indications of non-strict geographic distribution for *P. copri* groups

Although phylogenomics revealed the biogeographic distribution of groups, a few exceptions could be found in Figure 2A (e.g., g6 strains from China). The results based on genome bins may not sufficiently represent the population-level composition in each fecal sample because some strains could be missed during genome binning due to their low abundance or microdiversity (43). Two investigations based on a selected intra-species marker gene, *orth10* (See TEXT S1: Figure S3 for the reason to use this gene), were conducted to further investigate the strictness of the phylogeography.

The first study quantitatively assigned the metagenomic reads of *orth10* into groups. Analysis was conducted for 47, 70, 139, and 14 metagenomic datasets selected from Africans, Chinese, Europeans, and Americans, respectively, in accordance with *P. copri* abundance determined by its *gyrB* abundance (>10^-6^) in 1,267 integrated gene catalog database of human (IGC) samples and 67 African samples. All the four datasets showed a non-strict group-level distribution pattern, while the dominant groups were consistent with aforementioned biogeographic pattern (Figure 4A). Noticeably, the presence of g0, g1 and g7 in Europeans and detection of g0 in Chinese was revealed by this approach, was completely missing based on genomic information.

**Figure 4.**
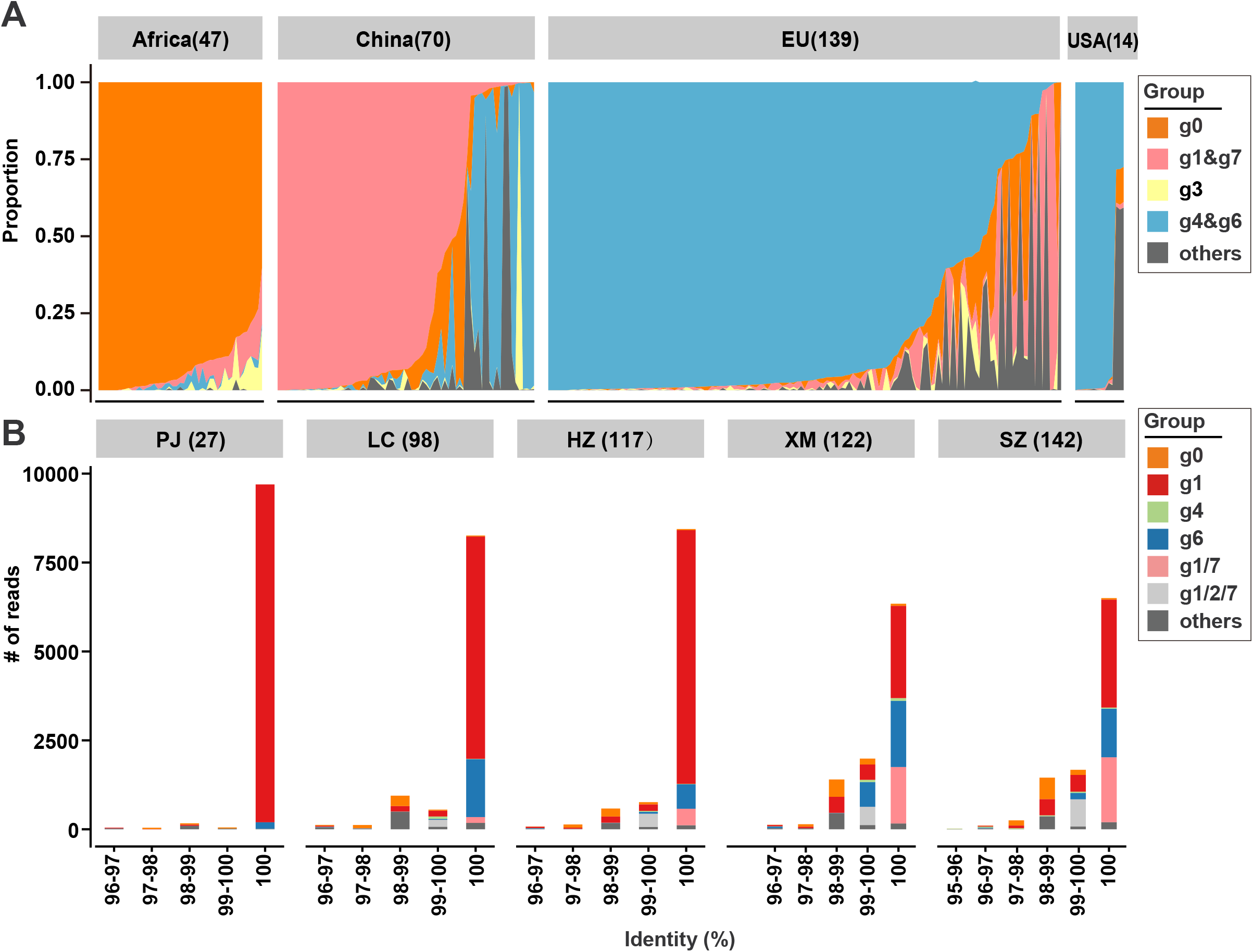
Non-strict biogeographical distribution for *P. copri* groups. A) Group-level profile of *P. copri* based on the full-length *orth10* in human gut metagenomes. The sample number with *P. copri* abundance >10^−6^ are shown in the brackets. Groups (i.e., g1/g7 and g4/g6) from the same geographical origin were merged to display. B) *P. copri* group-level profile in sewages of five cities in China based on the amplicon sequencing of the *orth10*. All samples are normalized to 10,000 sequences. In some cases, the short amplicon could not be clearly classified because of multiple top hits belonging to different groups (e.g. g1/g7). The number in brackets indicates the number of unique phylotypes detected in the sewage sample.

The second process was to conduct high-throughput sequencing for the *orth10* amplicons of sewage samples collected from five cities in China. As shown in Figure 4B, each sample contained 27–142 unique *orth10* phylotypes. Groups other than g1 and g7 (especially g0 and g6) were detected in the sewage samples, although sequences affiliated with g1 and g7 usually exhibited dominance. Although foreigners live in the cities, their contribution to the sewage was improbably high enough to change the main profile. Moreover, this result also suggested that the genome bins were a good representation of group-level diversity because all detected phylotypes are highly similar with the references (>95% and mostly 100% Figure 4B, TEXT S1: Figure S3 showed the similarity of almost all intragroup *orth10* exceeded 95% except group g0). Basing on the above results, we conclude that the group-level distribution in *P. copri* is not geographically strict, at least for the major groups.

### Long-term co-evolution and sporadic co-speciation of PCL with higher primates

Phylogeny based on 16S rRNA suggested that multiple PCL species have co-evolved with primates (Figure 1). Thus, the PCL abundance in fecal metagenomes (*n*=168) from 20 species of wild primates was analyzed based on the abundance of *gyrB* affiliated with PCL in these metagenomes (Figure 5A). Metagenomic assembly initially generated 82 PCL *gyrB* sequences, which represented 39 species-level clusters under 98% similarity cut-off (TEXT S1: Figure S4 shows the reason for using this criterion as the species-level cut-off in PCL). All these sequences were retrieved from eight species of higher wild primates (all from Cercopithecidae and Hominidae). The 39 representatives and 9 de-replicated (under 98% similarity cut-off) human gut *gyrB* sequences (extracted from the IGC database and isolates) were verified as PCL members because they formed a well-supported clade within *Prevotella* (Figure 5B, full-view in TEXT S1: Figure S5). Read-based quantification confirmed the absence of PCL in the guts of all lower wild primates and *Colobus guereza* (Figure 5A). The high diversity of previously unrecognized PCL species in higher primates strongly supported a long-term co-evolutionary history between the lineage and the hosts.

**Figure 5.**
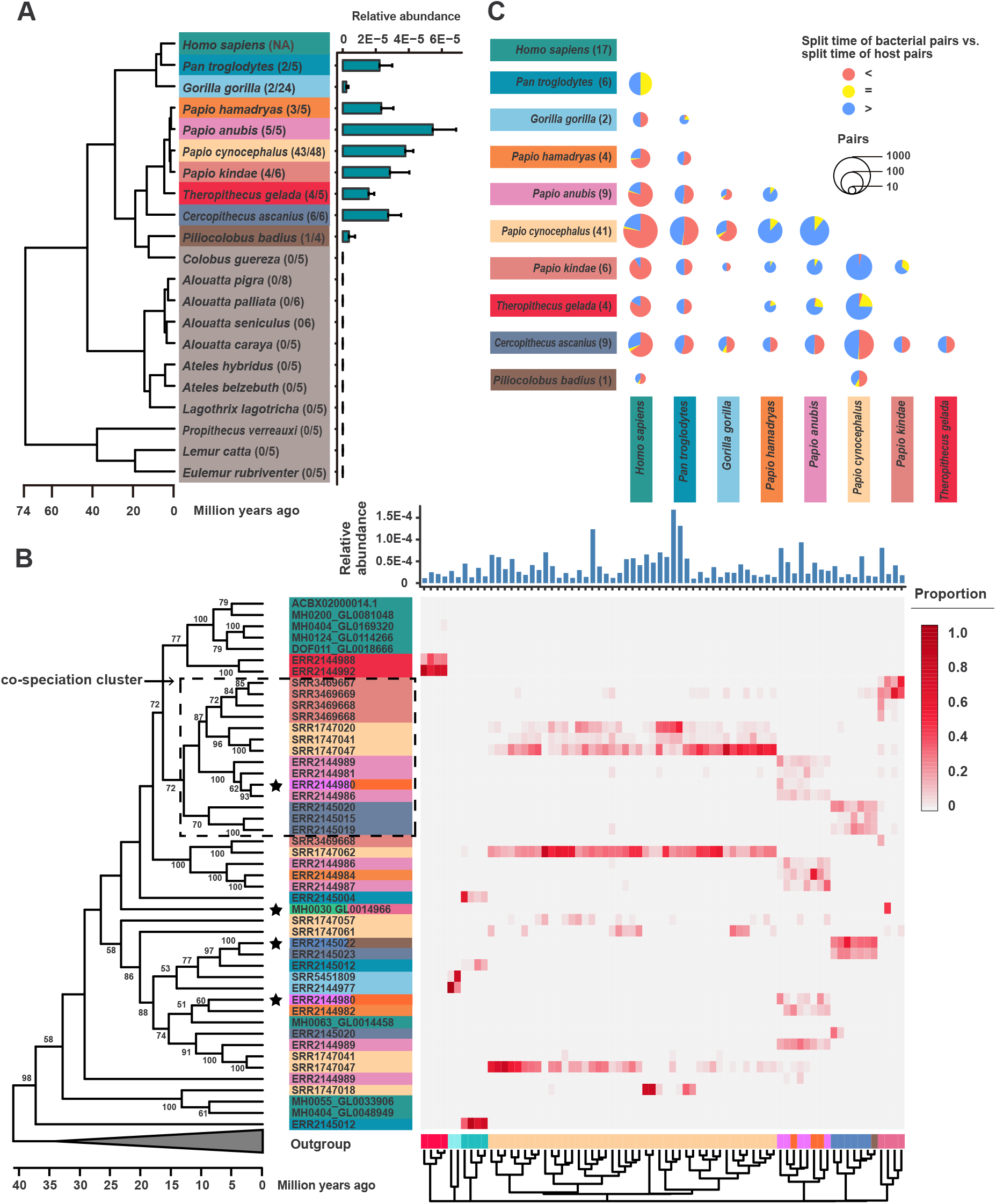
Diverse PCL members detected in the gut microbiome of nonhuman primates. A) Time-tree of hosts based on the evolutionary timescale. The relative abundance of total PCL in each host is shown in the barplot. B) Time constraint phylogenetic tree based on the 47 PCL *gyrB* representatives retrieved from wild nonhuman primates and human. The timescale is estimated by calibration based on the co-speciation cluster. Four representative sequences recovered from multiple host origins are marked by star. Read-based abundances are shown in the heatmap. Only samples with over 10^−5^ relative abundance of PCL in the metagenomes are listed with clustering according to Bray-Curtis dissimilarity (bottom). C) Pie chart showing the consistency between the split time of bacterial pairs and split time of host pairs. Only bacterial pairs more than 10 are shown. The number of *gyrB* sequences retrieved from the corresponding host are shown in the brackets.

Most host species were inhabited by multiple PCL species (Figure 5B). The PCL profile exhibited a conspicuous host-specific pattern, and no strong signals of phylosymbiosis were observed (clustering on the bottom in Figure 5B). Four *gyrB* representatives contained assembled sequences from multiple hosts, indicating their non-strict host specificity (Figure 5B). Phylogeny of *gyrB* hinted co-speciation events among four host species (*Papio anubis*, *Papio cynocephalus*, *Papio kindae*, and *Cercopithecus Ascanius*) with the furthest split time of 16.2–22.4 million years ago (Mya) (Figure 5A) but no signal across all higher primate hosts (Figure 5B). The split times of the four corresponding *gyrB* clusters (determined as the split time between hosts) were used as a reference to calculate the molecular clock rate and perform dating for the whole phylogenetic tree (Figure 5B). The initial date of PCL from other *Prevotella* species was deduced as 8.7–43.8 Mya (TEXT S1: Figure S5), which was highly variable but covered the split time of higher primates from others (28.0–31.4 Mya). Consistency was compared between the split times of hosts and bacteria (Figure 5C). Although the molecular clock rate for bacteria was highly variable, the two split times were still inconsistent in most pairs, except for some closely related host species such as between *Papio* spp. and between *Homo sapiens* and *Pan troglodytes*.

### Evidence of gene HGT among PCL members detected in the same host

In addition to *gyrB*, 16 PCL genome bins were retrieved from the fecal metagenomes of nonhuman primates (only three host species, i.e., *Papio cynocephalus, Pan troglodytes*, and *Gorilla gorilla*). Phylogeny of these strains representing seven uncultured species (designated as s1–s7 according to ANI values), *P. copri*, and Y7XP/Y7FG based on concatenated universal genes was generally consistent with that based on *gyrB* (Figure 6A and TEXT S1: Figure S6). The CAZys of the seven species highly overlapped with those of *P. copri*. The few absent CAZys in genomes of the *P. copri* include CE3, GH30_2, GH39, and GH76 that putatively target xylan or mannan (Figure 6A), which are important plant cell wall components (44). These polysaccharides are reasonably less abundant in the diet of modern human beings than in the diet of wild primates.

**Figure 6.**
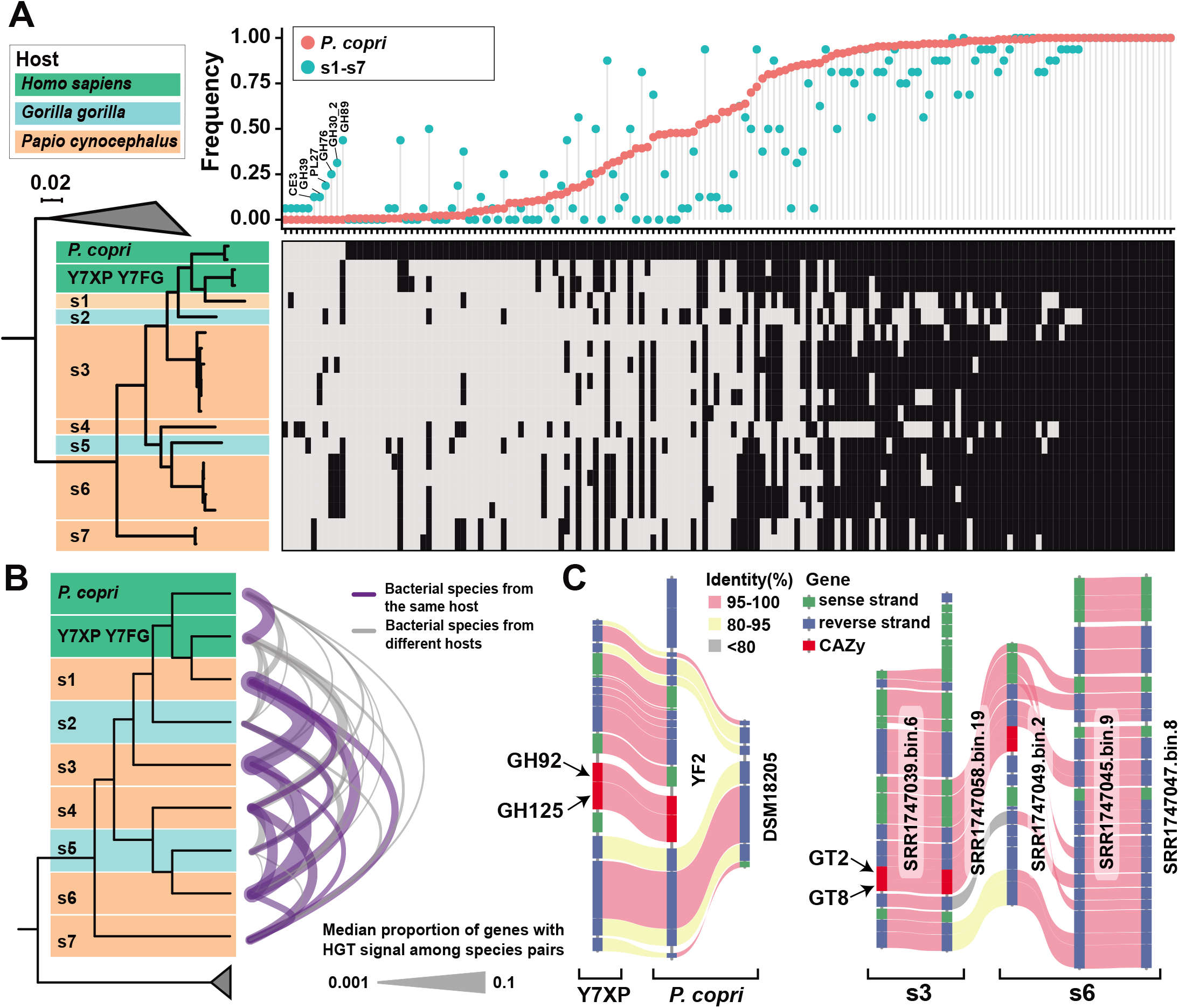
CAZys and HGT events of genomes affiliated within PCL. A) Phylogenetic tree of genomes affiliated within PCL inferred using 120 universal marker genes under the JTT model (left). The background color indicates the host origin. CAZy families are shown in the heatmap (black, present; grey, absent). The dropline plot shows the frequency of the CAZy families in 130 *P. copri* genomes (red) or 16 nonhuman primate derived PCL genomes (blue). B) HGT events among species of PCL. C) Two examples of selected CAZy genes with HGT signal shown by genomic synteny.

Since the HGT signal for CAZy genes has been detected among sympatric *P. copri* groups (Figure 3C), the potential gene HGT events among PCL members were examined. The upper boundary of 95% confidence interval of universal marker genes was set as the threshold for recognizing the HGT genes (TEXT S1: Figure S7). The bacterial species detected in different hosts only shared 0.6% genes with HGT signal, while the value is 4.0% for species from the same host (median value, Wilcoxon test, *P*<2.2×10^−16^, Figure 6B). Although this phenomenon may be partially attributed to the potential genomic contamination for metagenome-derived genomes, analysis based on isolates still showed the high proportion of HGT signals for the species from the same host (the median proportion between 24 *P. copri* isolates and Y7XP/Y7FG is 4.3%). Figure 6C shows the gene synteny of representative CAZy genes with HGT signal, which were putatively occurred in homologous genomic regions.

### PCL members in captive mammalian hosts were recently gained from humans

On the basis of the above results, PCL was hypothesized to have co-evolved with higher primates for a long period. However, *P. copri* and related taxa were widely detected in diverse non-primate captive mammals. Whether these taxa have evolved vertically or horizontally transferred to non-primate mammalian hosts remains unknown. Therefore, potential PCL *gyrB* sequences were extracted from the gene catalog of pigs and mice. Six pig-derived PCL *gyrB* sequences were phylogenetically affiliated with PCL, but no mouse-derived PCL *gyrB* was found (Figure 7A). However, three of the six pig-derived sequences were clustered (>98% similarity) with the human-derived *gyrB* representatives extracted from IGC database. The other three sequences still shared >95% identity to human-derived *gyrB* representatives. Noticeably, de-redundancy at a cut-off of 95% was conducted for the genes in the IGC database (45). Hence, every pig-derived PCL member has close relatives in human-derived members, but not *vice versa*. This finding suggests that these PCL species are horizontally transferred from humans.

**Figure 7.**
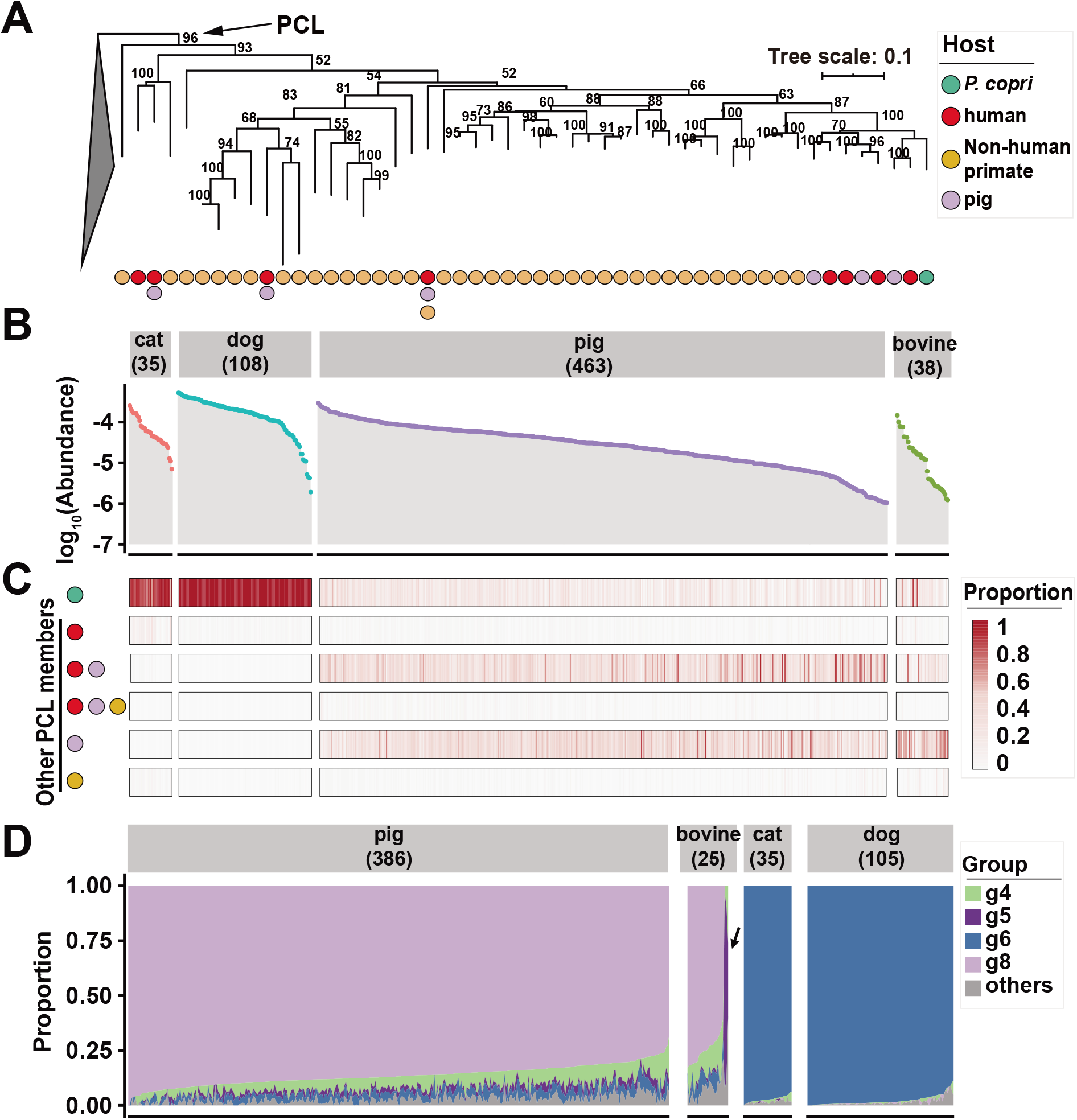
Distribution of PCL members and group-level profile of *P. copri* in the captive mammals. A) Phylogenetic tree based on representative *gyrB* sequences affiliated within PCL retrieved from humans, nonhuman primates, and pigs. The circle color shows the host origin. B) Relative abundance of total PCL in mammalian gut metagenomes according to their hits against the *gyrB* database. Only samples with the abundance of total PCL *gyrB* >10^−6^ are displayed and the numbers of samples are presented in the brackets. C) Heatmap shows the proportion of PCL members retrieved from different host origin in the captive mammals. D) Group-level profile of *P. copri* in the mammalian gut metagenomes according to their hits against the *orth10* database. The number shown in the brackets represents the samples number of animals with *orth10* abundance >10^−6^.

In reference to the above *gyrB* sequences, PCL was detected in fecal metagenomes from cats (*n*=36), dogs (*n*=125), pigs (*n*=533), and bovines (*n*=52). Figure 7B shows that the total PCL in the samples had an abundance over 10^−6^ (one assigned as PCL *gyrB* per million reads, requiring >95% similarity and 90% coverage, see TEXT S1: Figure S8 for the reason of the similarity criterion). In particular, 35, 108, 463, and 38 samples passed the abundance threshold for cat, dog, pig, and bovine, respectively. All these hits were profiled into six catalogues including *P. copri*, human-derived, human-pig shared, human-pig-primate shared, pig-derived, and primate-derived PCL members other than *P. copri* (Figure 7C). The PCL species detected in cat and dog samples were dominated by *P. copri*, and those in pig and bovine samples were mainly inhabited by *P. copri*, human-pig shared species, and pig-derived species.

Considering that *P. copri* was widely detected in these samples, group-level profiles of *P. copri* were established in the pig (*n*=386), bovine (*n*=25), cat (*n*=35) and dog (*n*=105) samples with high *orth10* (>10^−6^) abundance. As shown in Figure 7D, *orth10*-based quantification showed that cats and dogs (sampled from Europe and North America, respectively) almost exclusively harbored g6 strains, which geographically co-occurred in European and North American hosts. Pigs and bovines from Asia and Europe and bovines from North America were all dominated by g8. Bovines from Salvador were dominated by g5, a group also dominating the gut of Salvadorians.

## Discussion

### Co-evolutionary history for PCL in higher primates

Microbial samples from wild animals instead of captive ones are fundamental in determining their phylosymbiosis and co-evolutionary relationships to minimize artificial influence from humans (23, 46, 47). Besides that, our work emphasizes the need to use a comprehensive data source from multiple hosts in diverse geographical regions to obtain panoramic information (15, 29, 30). *De novo* retrieval of PCL genomes from the metagenomes of most human and animal samples must be conducted due to the high genomic divergence and microdiversity. In addition to genomes, the most comprehensive and accessible biomarkers rRNA and *gyrB* gene sequences (although not the most precise) can also be used as references to bypass the limitations of genome binning (e.g., low abundance and microdiversity).

A recent study found that *P. copri* complex in human gut comprises four species-level clades based on the genomes retrieved from metagenomes via referring to a few core-genomes (30). The current work discovered that PCL members in the human gut and higher primates are far more diverse than only four species according to *gyrB* sequences and genomes from expanded host spectrum. Similar to the co-speciation of *Bacteroide* spp. detected in extant hominid species (10), a signal of a few PCL members was found in four higher primates. However, the overall phylogenetic inconsistency suggested extensive horizontal transfer and extinction for the PCL members. A recent study showed the strong influence of environmental microbes on the gut microbiome of baboons (48), thus representing a possible pathway of host jumping. Our data of sharing the same PCL species amongst different wild hosts provided additional evidence for recent host jumping (Figure 5A). In addition, the extinction of certain species, which may be related to diet and behavior change as observed in experimental animals over several generations (49), may also play important roles in the distribution of PCL members.

### Are gut bacterial species shared by remotely related hosts evolved independently?

Genomic analysis further confirmed the intraspecific diversity and biogeographical pattern of *P. copri* (15, 29). Different *P. copri* groups shared critical or lower values to the species-level ANI and dDDH, suggesting their rapid evolutionary rates, which has also been proposed for endosymbionts (50), and experiencing allopatric speciation, a major mechanism for bacterial speciation (51). Coincident with *H. pylori* (37) and *Eubacterium rectale* (52), our results indicate that *P. copri* has a consistent phylogeographical pattern with human migration history, thus allowing calibration for its genomic evolutionary rates. As the low ANI values between conspecific strains in *H. pylori* have also been reported (53), the high evolutionary rate in *H. pylori* and *P. copri* raises a concern on whether a certain symbiotic bacterial species or clade shared by remotely related hosts has thoroughly evolved across the host phylogeny. Several previously reported gut bacteria, such as *Lactobacillus reuteri*, *Enterococcus faecalis*, and *Enterococcus faecium*, have experienced host-driven evolution across a wide range of mammalians or even vertebrates (12, 14). Oh et al. (12) deduced the split time of *L. reuteri* strains from multiple hosts by referring to a low mutation rate (54), resulting in molecular dating approximately 10 Mya. However, the low mutation rate has been suspected to be caused by the outdated methodology (55). If these species evolved at rates comparable with those of *H. pylori* and *P. copri*, then they are not likely to continuously co-evolve with the hosts for a long period without speciation. The conspecific ancestral strains were possibly incorporated into the gut microbiome of various hosts and have recently initiated a host-driven adaption (21). Comprehensive surveys on various symbiotic bacterial species across hosts will provide convincing evidence on their diversifying processes.

### Non-strict biogeography and host for PCL members suggest limited transmission barrier and potential niche selection

Poor host and geographical barrier have been observed in animal-associated microbial transmission (21, 47, 56, 57). Non-strict biogeographic distribution for subspecies-level profile of *P. copri* in human gut was proven by our study as well as by Tett et al. (30). The current work also revealed the putative extensive transmission of PCL members within different higher primates. Different (or at least for some) groups of *P. copri* and PCL members are distributed more ubiquitously than expected to a large extent following the microbiological tenet “everything is everywhere, but, the environment selects” (58). Factors other than geographical and host isolation, such as host diets and behaviors, and environmental characteristics in a given location have favored their occurrence and dissemination in the local population and certain host species. This explanation could be supported by the strain-level profile of *P. copri* being associated with different habitual diets (59); and some group-specific CAZys (e.g., alginase in g1 and hyaluronidase in g4/6) detected in our study.

Although no PCL *gyrB* sequence was detected in mouse gut bacterial gene catalog, which was generated from common laboratory mice (non-humanized) (60), a recent study verified the reliable transmission of *Prevotellaceae* from human feces to germ-free mice (61). Our investigation on the PCL members in captive mammals suggested that these bacteria recently originated from sympatric human hosts and potentially experienced niche selection, e.g., g8 of *P. copri* in bovines and pigs. Group g8 might be selected by the farming mode (e.g., diet) for the bovines and pigs. The bovines in Salvador, which were dominated by g5, were domesticated in different ways (e.g., the animals may be closer to humans than the industrial farming mode and fed with different diets) (62). Further comparison on the metabolic features of these animal-derived strains and human-derived strains may illustrate the evolutionary shaping of host adaptation within a limited period.

### Potential relevance of intra-lineage horizontal transfer of functional genes

Each host provides a distinct niche (or a collection of sub-niches) that can be colonized by bacteria (21, 63). Horizontal gene transfer and recombination are the main drivers of genomic divergence (64, 65) and play key roles in ecological adaption to new niches [83]. HGT is facilitated by the closely related phylogeny of donor and acceptor (66). Despite focusing on the genes encoding CAZy and uncharacterized mechanisms, our results provided evidence that HGT events have widely occurred among sympatric conspecific strains and closely related intra-lineage PCL members from the same host. The unique glycan degradation capability is important for gut colonization and sustention in human gut bacteria (67). Niche-driven rapid gain and loss of these genes within a large exchangeable pool may render the PCL members to be highly superior in the source competition. Moreover, extensive intra-lineage HGT events may result in the unreliable determination of specific phenotypes by group-level (or subspecies-level) and species-level identification. Caution must be adopted when linking phenotypes with taxa because the novel gene function may be rendered by recent HGT. A direct comparison among positive and negative strains is preferred (27).

In summary, our study focused on the macroevolution and microevolution of PCL in the guts of higher primates and humans. The results provided panoramic insights into the multiple effects of vertical transfer, horizontal transmission, and niche selection on the host and biogeographical distribution of a certain gut bacterial lineage. Studying the effects of PCL or other co-evolutionary lineages in animal guts on host phenotypes (e.g., health or disease) from the co-evolution perspective can aid the understanding on the interactions between host and gut microbes.

## Materials and methods

### Data collection for 16S rRNA gene, *gyrB*, metagenomes, and genomes

16S rRNA gene sequences from type strains and clones classified as *Prevotella* (*n*=534) were downloaded from EzBioCloud (68). Seven additional sequences were obtained from *P. copri*-like isolates (GCF_002224675.1, GCA_001405915.1, and five contributed by this study). SILVA SSU reference database (version 132) was used to track the host origins of *P. copri*-related sequences (69) (see TEXT S1: SI methods for the details).

As a species-level marker, the *gyrB* sequences of *Prevotellaceae* members in human gut were retrieved from the integrated gene catalog database of human, pig and mouse gut microbiomes (45, 60, 70), the metagenomic assemblies of wild nonhuman primates and 50 reference genomes (see TEXT S1: SI methods for the details). The *gyrB* sequences affiliated within PCL were included in the database to profile PCL members in the gut metagenomes of humans, nonhuman primates, and captive mammalian hosts (see TEXT S1: SI methods for the details).

A total of 2,784 publicly available gut metagenomes of humans, nonhuman primates and other mammals were collected from 21 studies involving 26 host species from 30 countries (DATA SET S1: Table S4). Forty-eight published *P. copri*-like genomes, including 21 isolates and 27 genome bins from African (30, 42), were collected. The present study contributed 119 new genomes (5 from isolates and 114 from metagenomes). Information for all genomes was listed in DATA SET S1: Table S1.

### Isolates and genome sequencing

Fresh stool samples were collected from four healthy Chinese volunteers (previous investigation on their gut microbiota suggested high abundance of *P. copri*-like taxa) and immediately transferred to an anaerobic glovebox (N_2_: CO_2_: H_2_=80: 15: 5) for isolation on YCFA medium (71). Colonies were picked after cultivation at 37 °C for 96 h. Full-length 16S rRNA gene sequences of the isolates were used to identify *P. copri* and its related strains on the EzBioCloud platform (68).

Genomic DNA of *P. copri* and their related species isolated in this study was extracted and sequenced with PE150 strategy on Illumina Hiseq 4000 platform (commercial service, Novogene, Beijing). *De novo* assembly was performed by SPAdes v3.9.0 (72). Only scaffolds longer than 1,000 bp were included in the downstream analysis. The whole genome of YF2 strain was achieved by combining Illumina and PacBio RSII platform sequencing (commercial service, Novogene, Beijing).

### Genome binning, quality assessment, and annotation

Genomic binning using mmgenome was manually performed to obtain high-quality *de novo* assembled genomes of *P. copri* and related taxa from humans and nonhuman primates, respectively (73). Prescreening of the 1,679 gut metagenomes from humans (DATA SET S1: Table S4) was conducted using the relative abundance of *P. copri* as estimated by the relative abundance of *gyrB* (usually >10^−5^ for IGC data) or MetaPhlAn v2.0 (>10% relative abundance for non-IGC data) to improve the efficiency (74). For the 168 gut metagenomes from nonhuman primates (DATA SET S1: Table S4), raw reads were quality filtered with Trimmomatic v.0.36 (75). The raw reads of selected human and nonhuman primate samples were first assembled using SPAdes v3.9.0 (72). Only scaffolds longer than 1,000 bp were retained for genome binning. The raw reads were mapped to the scaffolds using Bowtie2 v2.2.9 (76), and the coverage profile was calculated by SAMtools v0.9.1 (77). Other necessary files were generated using script data.generation.2.1.0.sh (73).

Completeness and contamination of all draft genomes were assessed by CheckM v1.0.7 (78). Pairwise ANI and dDDH values among *P. copri* genomes were calculated by FastANI v1.3 (79) and Genome-to-Genome Distance Calculator 2.1 (35), respectively. Genes encoding CAZy families were annotated using HMMER 3.1b2 (80) against dbCAN HMMs v6 (81), and the results were filtered according to the recommended threshold.

### Defining core protein orthologues of *P. copri*

Core orthologous gene clusters of *P. copri* genomes were defined using the method of Oyserman et al. (82) with modifications. All incomplete open reading frames (ORFs) with potential redundancy (i.e., multiple fragments from one ORF) were cleaned prior to downstream analysis (see TEXT S1: SI methods for the details). All-against-all BLASTP was performed for cleaned ORFs from *P. copri* genomes (83). Identity and inflation values were determined according to McCill et al. (84) by maximizing the maintenance of genes with the same function in a cluster. Orthologous gene clusters were generated by MCL with optimized inflation value of 1.2 (85). A total of 1,095 single-copy core orthologs that appeared in more than 90% of the *P. copri* genomes were determined.

### Phylogenetic analysis

Phylogenetic analyses were performed for single genes and concatenated alignments of single-copy core ORFs. The trees based on single genes were reconstructed using MEGA v6.06 with 100 bootstrap iterations (86), and those based on concatenated genes were reconstructed by maximum likelihood analysis using RAxML v.8.2.4 (87) or FastTree v2.1 (88) on CIPRES web server (89). A truly whole-genome-based phylogenetic analysis of the coding sequences was conducted at the nucleotide level using the latest version of the Genome BLAST Distance Phylogeny method under recommended settings (35, 90). All phylogenetic trees were visualized via the iTOL web server (91). Further details are described in TEXT S1: SI methods.

### Determination and application of a quantitative gene with intraspecific resolution for metagenomes

A quantitative marker gene with intraspecific resolution must be selected because *P. copri* has high genomic diversity that may cause quantitative biases at sub-species level in metagenomes. For *P. copri* genomic pairs, the best candidate was determined by calculating the Spearman correlation of distances between concatenated 1,095 single-copy core orthologs and each single core ortholog. The optimized gene was designated as *orth10* (the corresponding gene of type strain DSM 18205 was EFB36125.1, a response regulator receiver domain protein) and was used as the basis for the group-level profiling of *P. copri* in human and mammalian gut metagenomes and the investigation on *P. copri* populations in raw sewages collected from five cities of China (See TEXT S1: SI methods for details).

### Molecular dating for the split times between *P. copri* groups and between PCL members

Molecular dating was conducted as previously reported by Oh et al. (12). PHI test was used to identify the intragenic recombination of 120 universal genes as proposed by Parks et al. (92, 93). The dN/dS ratios for genes without significant intragenic recombination (PHI test, *P* >0.05) were calculated by using KaKs Calculator (94). Split time was estimated by the synonymous mutation rate among various groups and the long-term mutation rate of housekeeping genes of another human gut symbiont, namely, *H. pylori* (2.6×10^−7^ per site per year) (36).

The divergence time of the *gyrB* sequences retrieved from IGC and nonhuman primates was estimated by Bayesian MCMC analysis implemented in BEAST2 v2.5.2 (95). The bacterial lineages showing signals of co-speciation with primate hosts were used as calibration, and the maximum likelihood tree inferred by MEGA was employed as the starting tree. The analysis was run 50 million generations and sampled every 1,000 steps under the GTR+G+I substitution model with a lognormal relaxed molecular clock (10). Tracer v1.7.1 (http://tree.bio.ed.ac.uk/software/tracer/) was utilized to ensure that the effective sample size was larger than 200 for all parameters. The tree files were summarized in TreeAnnotator with the first 25% discarded as burn-in (96).

### Determination of HGT events among PCL species

All-against-all BLSATN was preformed between heterospecific genome pairs to define the HGT events among PCL species. Shared genes with high similarity between any two heterospecific genomes were classified as HGT due to the lack of available tools to identify HGT events among closely related species. For a given species pair, HGT signal threshold was set as the upper boundary of the 95% confidence interval of similarity between complete universal genes (92). Gene pairs with similarity higher than the threshold were recognized as HGT positive, and the proportion of genes with HGT signal was calculated for each genome pair 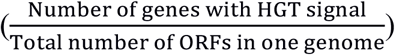.

### Statistical analysis and visualization

Statistical analysis was conducted in R 3.5.1. The *rcompanion* (97) package was used for Fisher exact tests, and *ggplot2* (98), *pheatmap* (99), and *ggalluvial* (100) packages were applied for data visualization. The genomic synteny of the fragment containing CAZy was visualized by MCscan (Python version) (101).

## Acknowledgements

Prof. Xin Yu and Mr. Chengsong Ye are thanked for their help in obtaining the sewage samples.

## Funding

This study was supported by the National Natural Science Foundation of China (No. 31670492 and No. 31500100).

## Data availability

The 16S rRNA gene sequences contributed by this study have been deposited in Genbank under accession numbers MN658562-MN658566. The genomes recovered in this study have been deposited in the Sequence Read Archive (SRA) under accession number PRJNA555508. The genome sequencing data of isolates in this study and the *orth10* amplicon sequencing data had been deposited in the SRA under accession number PRJNA555745, PRJNA565808 and PRJNA557417.

## Author contributions

FG conceived and supervised the study. HL and FG designed the study. CXH and JZ performed the experiments. HL, JMK, ZJW and FG analysed the data. HL and FG wrote the manuscripts. JMK, WZ and YT reviewed and provided valuable edits to the manuscript.

## Ethics approval and consent to participate

Not applicable.

## Consent for publication

Not applicable.

## Competing interests

The authors declare that they have no competing interests.

